# Stimulus-Specific and Generalized Taste Aversion Behaviors and Their Relationship to Cortical Dynamics

**DOI:** 10.64898/2026.06.02.729621

**Authors:** Christina T. Mazzio, Sydney Flashman, Abuzar Mahmood, Natasha Baas-Thomas, Jian-You Lin, Karolina Komar, Donald B. Katz

## Abstract

Aversive taste behaviors, such as gaping, are commonly viewed as fixed, hard-wired responses important for ejecting potentially toxic tastes from the mouth. Yet, taste responses are highly susceptible to modulation by experience and context; for instance, conditioned taste aversion (CTA), a form of learning in which rats are made to respond aversively to a sweet taste after it has been paired with gastric malaise, can cause gaping to previously acceptable tastes. Here, we compare aversive responses to saccharin sodium (sacc CS) caused by repeated aversive experience (in the form of CTA training) to those caused by naturally aversive quinine HCl, and how these responses are aligned with gustatory cortical (GC) activity. Using simultaneous single-neuron ensemble recordings in GC and electromyography (EMG) recordings in the jaw-opener muscle in awake, freely-moving rats and a machine-learning based behavioral classifier, we not only identified two classes of gaping, but characterized how these classes reflect different mechanisms driving aversive behavioral outputs. The first gape class is taste-specific and emerges following GC ensemble dynamics associated with taste identity and palatability coding, leading to canonical gaping responses to naturally or learned-aversive stimuli. The second is rapid-onset (<200 ms following taste delivery), internal state-dependent gaping the animal exhibits to all tastes in a state of heightened vigilance. This generalized aversion is subsequently suppressed once palatability-related responses emerge in GC, such that only gapes to truly aversive tastes remain. Together, these findings demonstrate that gaping is not a unitary reflex but is instead driven by multiple aversion judgment systems active in different contexts.

## Introduction

A rat faced with an unappealing substance on the tongue responds with a set of behaviors that signal that taste’s aversiveness. Most notable among these is an oral behavior called the “gape,” a rhythmic, yawning jaw and tongue movement designed to facilitate ejection of the offending substance from the oral cavity (Grill & Norgren, 1978a; Steiner et al., 2001; Pereira et al., 2021); gaping minimally requires brainstem circuits (Grill & Norgren, 1978b), but *in situ* can be driven by gustatory cortical (GC) processing—specifically, by the sudden ensemble transition of firing from stimulus- to palatability-related that occurs approximately 1 sec after taste delivery (Sadacca et al., 2016; Mukherjee et al., 2019). Other behaviors—limb flails and chin rubs, for instance—emerge to reflect particularly strong aversion, sometimes replacing gapes (Grill & Norgren, 1978a).

While it is often assumed that these aversion-related behaviors are “hard-wired” reactions to bitter tastes, it has long been known that the palatability of all tastes is exquisitely sensitive to context and experience. A palatable taste that is paired with gastric malaise, for instance—a form of learning called conditioned taste aversion (CTA)—becomes deeply aversive (Garcia et al., 1955 & 1966), such that it may drive gaping (Berridge et al., 1981; Pelchat et al., 1983; Parker, 2003 & 2006). Even benign experience, including experience that isn’t taste related (Gutiérrez-Vera et al., 2022), can change rodent (De La Casa & Lubow, 1995; Flores et al., 2018) and human (Mennella et al. 2001; Prescott et al., 2002) taste preferences. Thus, lab rats differ with regard to what tastes they consider aversive (Maigler et al., 2026).

The fact that experience can have such profound impact on the driving of aversion motivates the hypothesis that aversive behaviors can be driven in multiple ways, and that the circuit involved in driving rejection can be different in different contexts. Most obviously, it is reasonable to suggest that CTA learning drives gapes that differ in some way from naïve gaping to quinine, a bitter tastant. This would not be surprising: many learned behaviors are distinct from their naïve counterparts—conditioned and naïve eyeblinks, for example, differ in timing, kinematics, and driving circuit; McCormick et al., 1981, 1982, & 1984; Steinmetz et al., 1987 & 1992); furthermore, the involvement of GC in learned gaping has been suggested to be distinct from its involvement in naive gaping to quinine (Kiefer & Orr, 1992). Simply put, there may be multiple circuits driving individual aversive behaviors, and driving aversion in general.

Here we investigated this possibility in rats given CTAs to saccharin, using simultaneous single-neuron and electromyographic (EMG) recordings in conjunction with recently developed machine-learning analyses. Our data revealed CTA-induced gaping to saccharin that differs from naïve gaping to quinine, but deeper analysis made it clear that these differences reflect the existence of two distinct mechanisms that can trigger aversive responding. Only one of these mechanisms is taste-related: while aversive responses can be triggered by a specific taste, rats anticipating the potential delivery of aversive tastes enter a state in which any stimulus in the mouth triggers aversion, and in which the natural aversiveness of quinine and established aversiveness of saccharin are enhanced. Furthermore, “state-dependent” (general) aversive responses occur early (within 0.2 sec of stimulus delivery; see also Halpern & Tapper, 1971), and are quenched by the very completion of taste processing dynamics that drives taste-specific gaping. While rats normally find fluids palatable until taste processing reveals otherwise, they can be driven into a state in which the opposite occurs.

## Materials and Methods

### Experimental Design and Data Collection

#### Subjects

Adult, female Long–Evans rats (n = 22; 17 CTA-trained, 5 sham-CTA) 250–300 g at time of electrode implantation, Charles River Laboratories) served as subjects in our study. The rats were housed in individual cages in a temperature- and humidity-controlled environment under a 12 h light/dark cycle and given *ad libitum* access to food (Labdiet 5p00 Prolab RMH3000) and water before the start of experimentation. To minimize taste preexposure, we ensured rats were housed singly, in clear plastic cages containing only sawdust (or paper post-surgery) bedding, and paper enrichment. All animals were weighed daily following surgery to ensure that they never dropped below 85% of their pre-surgery weight. All experimental methods were in compliance with National Institutes of Health guidelines and were approved in advance by the Brandeis University Institutional Animal Care and Use Committee.

#### Electrode, EMG, and intraoral cannula construction

Custom microwire bundle drives were made with 32 nichrome microwires per bundle (A-M Systems catalog # 761500). The microwire bundle was affixed to an electronic interface board (EIB) pre-soldered to a 64-ch Hirose connector (product # DF40C(2.0)-70DS-0.4V(51) (OpenEphys). All drives were implanted unilaterally, and therefore only half of the available channels on the EIB were wired in each drive. Two free pins on the EIB were wired to PFA-coated stainless-steel electromyography (EMG) electrodes (A-M Systems catalog #793200). The drive was grounded using two stainless steel wires. Design and construction details available at https://github.com/danielsvedberg/blech_shuttledrive_mk6.

Intraoral cannulae (IOC) were constructed from flexible polyethylene tubing (A-M Systems catalog #803100) and designed with a flanged tip and washer to ensure stability. After implantation, the IOC was connected to a plastic top complete with a locking mechanism to allow the delivery of tastants directly onto the tongue from inside the behavioral apparatus as described previously (Fontanini & Katz, 2006).

#### Surgical Implantation

All subjects were implanted with an IOC, EMG in the anterior digastric (AD) muscle, and a microelectrode array in left hemisphere GC. All rats were first anesthetized with an intraperitoneal injection of a ketamine/xylazine mixture (100 mg/kg and 5 mg/kg body weight, respectively), after which we shaved the scalp and a region of skin under the jaw that sits above the AD. The head was then situated in the stereotaxic frame and the scalp was cleaned with sterile ethanol and iodine wipes. After excising the scalp and leveling the skull, five self-tapping screws were drilled into the skull for supporting and grounding the electrode bundles. The ground wires on the microdrive were wrapped around a screw located over the cerebellum for grounding. For subjects that had an electrode implant, we next made an additional, larger craniotomy (∼2 mm dia.) above GC. An electrode microdrive (*see Electrode, EMG, and intraoral cannula construction*) was slowly lowered (over 5-10 min), and electrodes were positioned 0.5 mm above the GC (+1.4 mm A/P, +/- 5.0 mm M/L, 4.2 mm D/V). Afterwards, the craniotomy was sealed with Kwik-Sil (World Precision Instruments). The microdrive was secured in place with dental acrylic.

The rat was then removed from the stereotaxic frame and implanted with a single (right-side) IOC in the space between the first maxillary molar and the cheek, through the masseter muscle and inside the zygomatic arch, and out through the opening in the scalp (Katz et al., 2001; Phillips & Norgren, 1970) before being cemented in place. The EMG electrodes were channeled down the left side of the face (opposite from the IOC); after the overlying skin had been teased away from the belly of the digastric muscle, one end of each EMG electrode was then inserted into the muscle using a suture needle (for more details, see Dinardo & Travers, 1994; Li et al., 2016; Loeb & Gans, 1986; Travers & Norgren, 1986). The electrode wires were trimmed and held in place with Vetbond tissue adhesive (3M) and the skin covering the anterior digastric was sutured closed. Finally, bacitracin ointment was applied all around the base of the headcap and over the incision under the jaw. Rats were postoperatively injected with analgesic (Meloxicam 1-2 mg/kg), saline, and antibiotic (Baytril 5-10 mg/kg). Similar antibiotic, saline and analgesic injections were delivered 24, 48 and 72 hr later, and bacitracin ointment was reapplied to the wound at the same time points. The rats were allowed to recover for a minimum of 6 days before starting experimentation, and daily weight records were kept to ensure that the rats did not fall below 85% of pre-surgery weight.

#### CTA paradigm

Four days following surgery, the electrode bundle in the microdrive was driven down 0.25mm, and one day later, driven down another 0.25mm to the depth of the dysgranular layer of GC (-4.7mm). Animals were placed on a water-restriction regimen (15mL water/d), then habituated to the experimental environment for 2d, followed by 2d habituation to 60μl water deliveries through the IOC.

##### Experimental cohorts

All animals in the study contributed EMG data and fell into one of four experimental cohorts; trained, over-trained, combined test, and sham-trained. Trained animals received a single CTA training session, a testing session, and a quinine session, during which a naturally aversive taste, quinine hydrocholoride (QHCl), was administered (n = 7).

Over-trained animals received the same protocol as trained animals, except they received two identical training sessions (each separated by 24 hr) before the testing and quinine sessions (n = 4). The combined test animals received a single training session followed by one last testing session, during which the conditioned stimulus (CS) and QHCl were presented on the same day (n = 6). Sham-trained animals received the same protocol as the trained group, but the CS was paired with a saline injection during the training session so they would not acquire CTA (n = 5). See **Table 1** for details of each subject used in the study.

**Table 1.**
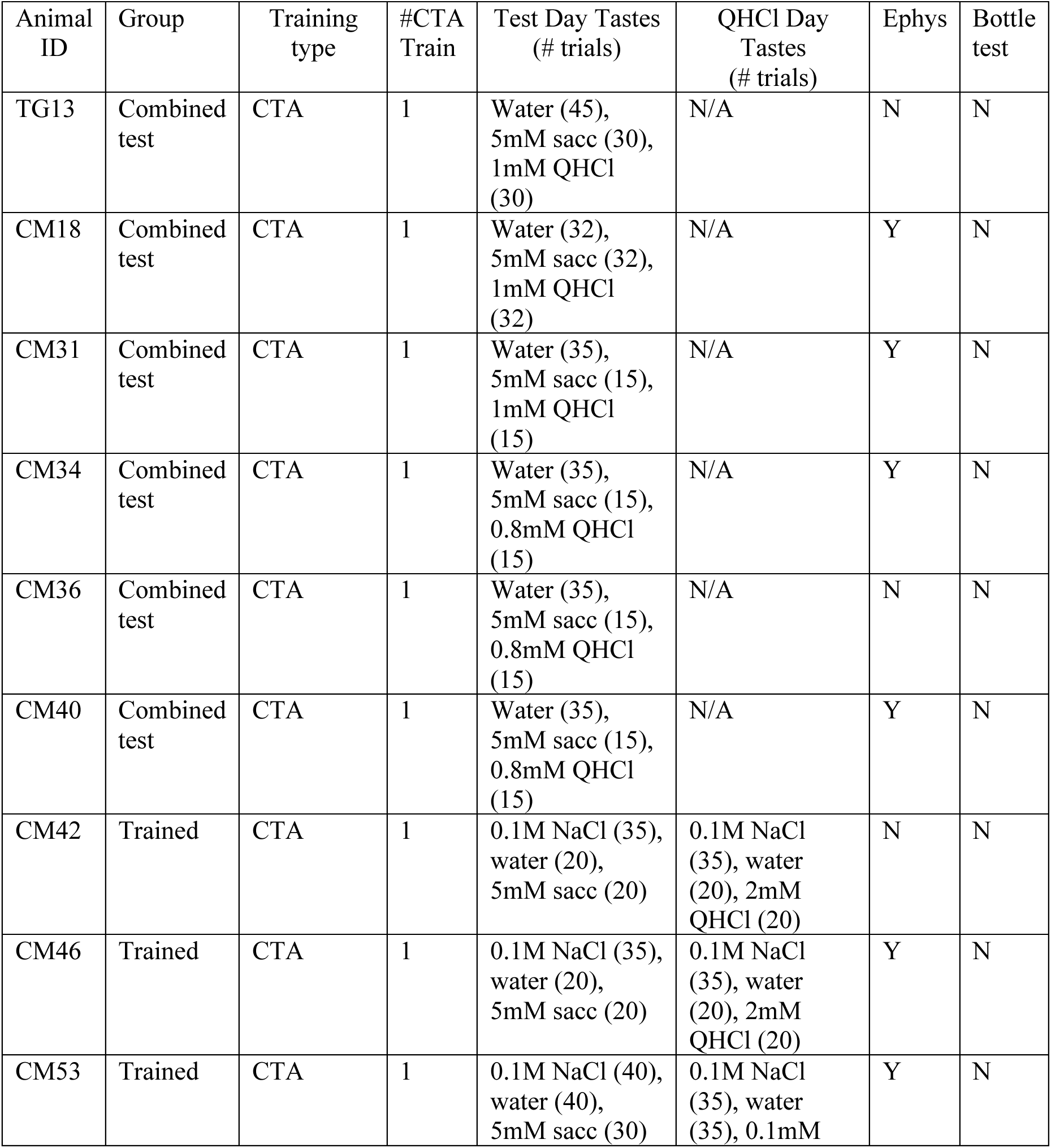
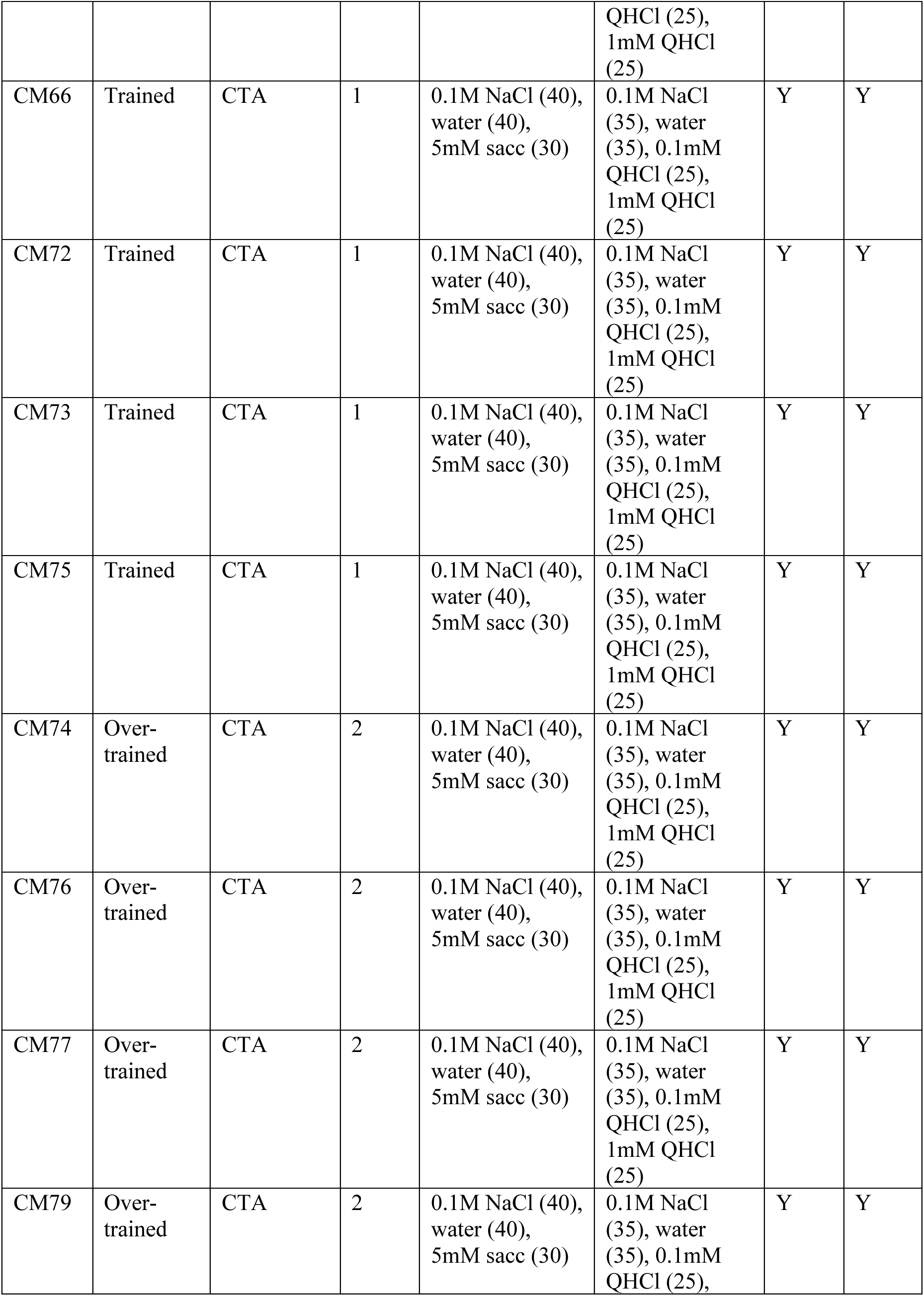
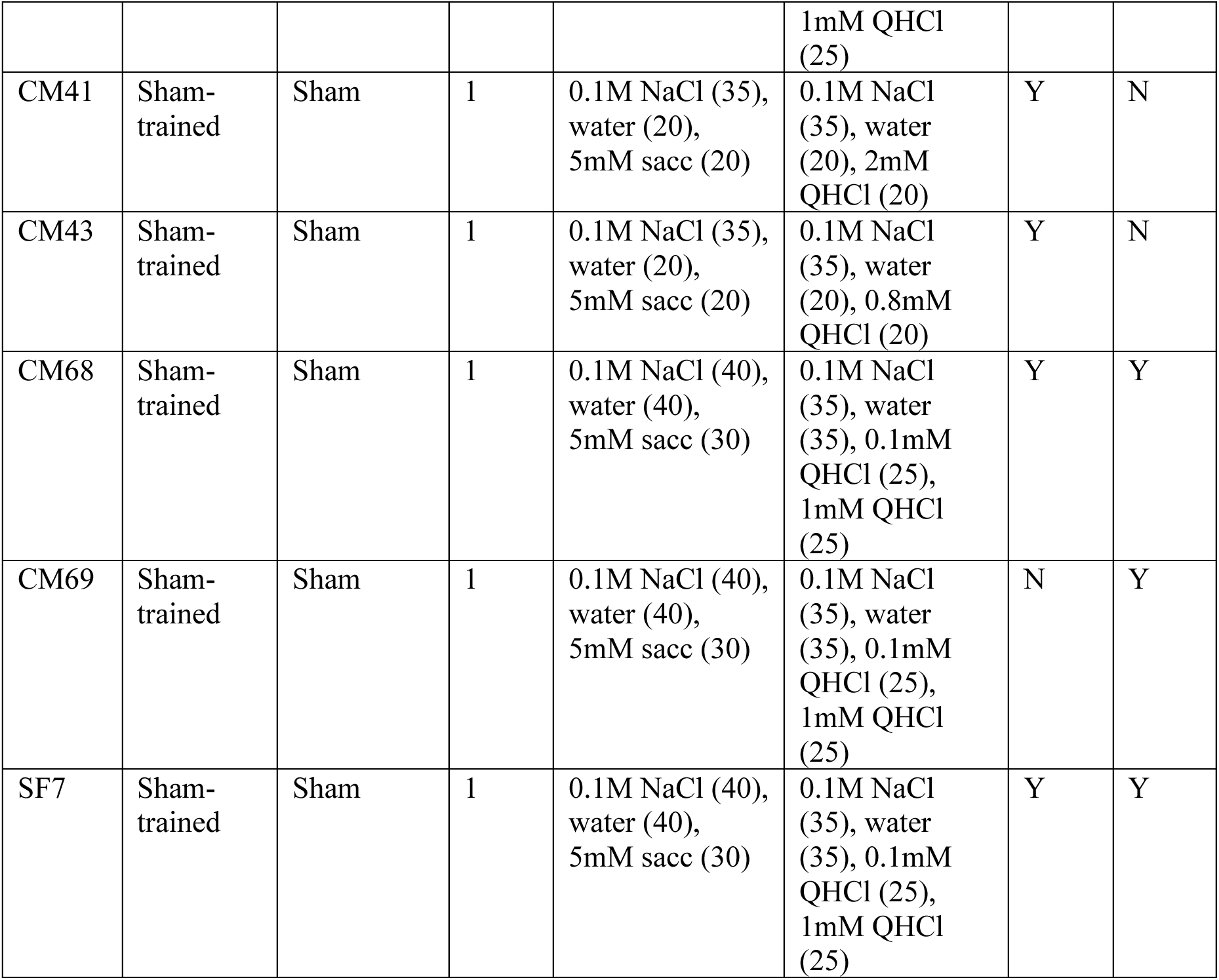
Summary of rat subjects and experimental conditions.

##### Training session

Following habituation sessions, animals underwent a CTA training session in which they received 60 deliveries of 60μl aliquots each of water and 5mM saccharin (sacc CS) through the IOC (120 deliveries total; 30s ITI). To prevent more than two consecutive deliveries of either taste solution, deliveries were made in blocks of two, with each taste delivered once within a single block. Rats remained inside the experimental rig for 30 min following the final taste delivery before receiving a subcutaneous injection of 0.6M LiCl (1% v/w) while inside of the rig. All animals remained inside of the rig for 45 min following injection to ensure they experienced the first 30 minutes of illness (starting ∼15 min after injection; Stone et al., 2019) in the same environmental context as the taste deliveries. Animals in the over-trained group received an identical training session 24 hr later. Rats in the sham-trained group followed the same protocol, except they received a subcutaneous injection of 0.6M NaCl (1% v/w). There was some trial count and QHCl concentration variability in the sham-trained and combined test groups (see **Table 1**).

##### Testing session

24 hr after the training session, animals were placed in the same experimental rig and given 10 deliveries each of 0.1M NaCl and water followed by 30 deliveries each of 0.1M NaCl, water, and sacc CS (110 deliveries total; randomized 25-35 intertrial interval, ITI). Deliveries were made in blocks of three, with each taste delivered once within a single block. QHCl was also given during this session to animals in the combined test group (see **Table 1**).

##### Quinine session

24 hr after the testing session, animals were placed in the same experimental rig and given 10 deliveries each of 0.1M NaCl and water followed by 25 deliveries each of 0.1M NaCl, water, 1mM QHCl, and 0.1mM QHCl (120 deliveries total; randomized 25-35 ITI). Deliveries were made in blocks of four, with each taste delivered once within a single block. Only animals in the trained, over-trained, and sham-trained groups had a quinine session.

##### Bottle Test

Due to variability in the amount of gaping to the CS between individual subjects in the same group, a subset of the trained, over-trained, and sham-trained groups underwent bottle testing to confirm CTA acquisition. To habituate animals to the bottle test, a bottle of 15ml water was placed in the home-cage 2-3hr after the last taste delivery, starting on the first water habituation day. The first day of the bottle test took place after the testing session; animals were given 30min access to 15ml sacc CS in a bottle placed inside the home-cage 2-3hr after the last IOC taste delivery and bottle weight was recorded before and after the test. Animals then received 30min access to a bottle of 15ml water in the home-cage immediately after. After the quinine session the next day, animals received an identical bottle test, but with access to 1mM QHCl during the first 30min bottle session. After all bottle sessions, animals were given access to the amount of water equal to the difference between 15ml and the volume consumed during the test, keeping daily maximum fluid consumption at 15ml in addition to ∼8ml given during IOC-delivery sessions.

#### Electrophysiology

Throughout each taste-exposure session, neural and EMG signals were simultaneously amplified and digitized by a 64-channel headstage (Intan, Open-Ephys) connected to the EIB, and recorded at 30kHz using an Intan RHD USB interface board (catalog# C3100) for offline analysis. In addition, timestamps of each stimulus delivery were recorded alongside electrophysiology. The experimental chamber was ensconced in a Faraday cage that shielded recordings from external electromagnetic influences.

#### Histology

In preparation for histology to confirm GC electrode placement, rats were deeply anesthetized with an overdose of the ketamine/xylazine mixture. We perfused the rats through the heart with 0.9% saline followed by 10% formalin and harvested the brain. The brain tissue was incubated in a fixing mixture of 30% sucrose and 10% formalin for several days before being sectioned into 50μm coronal slices on a sliding microtome (SM2010R, Leica Microsystems). Sections containing the electrode implant sites around GC were imaged with a fluorescence microscope (BX-Z710, Keyence). Histological verification for rats with only EMG implants was not necessary as targeting of EMG electrodes was directly observed during surgery.

### Data Processing and Statistical Analysis

The analysis of data and statistical tests were performed using custom-written software in Python as described below. Some of the more prominently used packages are mentioned here:

- Scipy (statistical analysis; Virtanen et al., 2020)
- Pandas (data handling and manipulation; The pandas development team, 2025)
- Scikit-Learn (machine learning; Pedregosa et al., 2011)
- Numpy (numerical computing; Harris et al., 2020)
- Blech_EMG_Classifier (electromyographical data processing; Baas-Thomas et al., 2026)
- Blech_Clust (electrophysiological data processing; Mahmood et al., 2025)
- Pytau (electrophysiological changepoint modelling; Mahmood et al. 2023)
- BlechCTA (behavioral and neural data analyses)

#### EMG processing

As detailed previously, we recorded voltage signals from two unipolar EMG electrodes implanted in the anterior digastric muscle at 30 kHz. We used the difference in the voltage recorded by the two electrodes as the EMG signal—this procedure helps to cancel out any large artifacts produced by the animal’s movements and is equivalent to using a differential amplifier (Li et al., 2016). We down-sampled the EMG signal to 1000Hz by averaging the voltage values in sets of 30 samples, and high pass filtered the down-sampled signal above 300Hz (Li et al., 2016; J. B. Travers & Norgren, 1986) using a 2nd order Butterworth filter. The absolute value/magnitude of the filtered EMG signal was then lowpass filtered (again using a Butterworth filter of order 2) below 15Hz, effectively capturing the envelope of variation of the EMG signal (plotted as a black curve in Figure 1C, right panel). This cutoff of 15Hz is sufficient for identifying orofacial behaviors, all of which occur at frequencies smaller than 10Hz (Li et al., 2016; J. B. Travers & Norgren, 1986).

**Figure 1.**
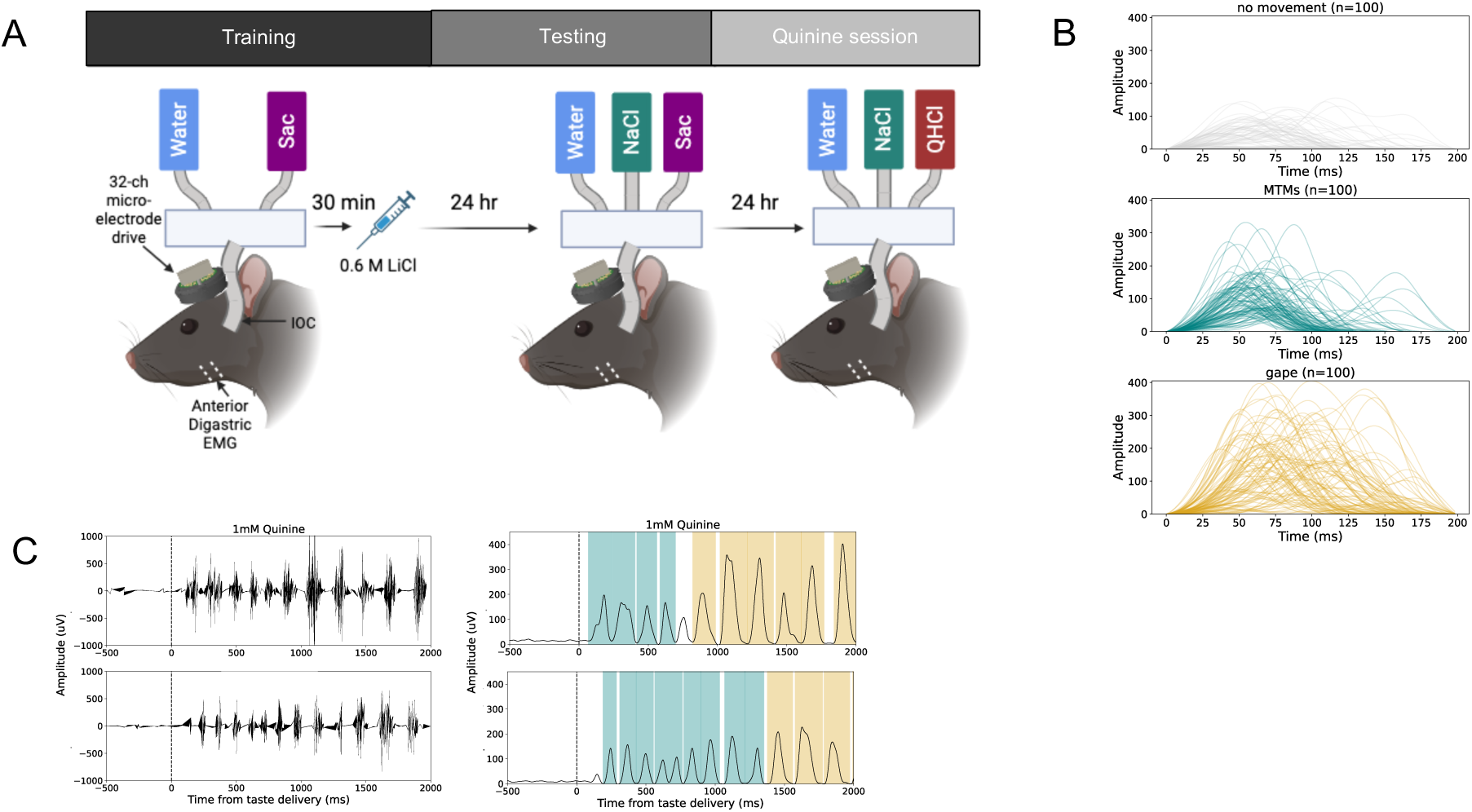
Rats produce naïve gapes to bitter quinine. **A)** Experimental design for 3-day CTA paradigm for the trained group. Animals are implanted electromyography (EMG) wires in the anterior digastric muscle and an intraoral cannula (IOC). Animals first undergo a training session in which deliveries of water and 5mM sacc followed by 0.6M LiCl (1% v/w) injection s.c. 30 minutes later. Sham-trained controls are injected with 0.6M NaCl (1% v/w) instead. 24 hr later, animals receive water, 5mM sacc, 0.1M NaCl, and water during a testing session. After another 24 hr, all animals receive 0.1mM QHCl, 1mM QHCl, water, and 0.1M NaCl during the quinine session. **B)** Overlain waveforms for each class of labelled movements detected by XGBoost EMG classifier. **C)** Example filtered (left) and enveloped (right) EMG signals recorded during two example quinine session 1mM QHCl trials. Gape (4-6 Hz) and MTMs (6.6-10 Hz) waveforms are highlighted in yellow and teal, respectively. Taste delivery at 0ms. Sacc, saccharin sodium; CS, conditioned stimulus; NaCl, sodium chloride; QHCl, quinine hydrochloride; LiCl, lithium chloride; MTMs, mouth movements.

#### XGBoost EMG classifier

We used a machine-learning classifier to detect and identify individual orofacial movements from anterior digastric EMG activity (Baas-Thomas et al., in press). This classification model is based on a gradient-boosted trees algorithm (XGBoost; Chen & Guestrin, 2016) and is trained on a suite of features from each waveform including amplitude, multiple frequency-based properties including maximum frequency, time from previous peak, time to following peak, and duration, and shape-based properties extracted using the first three principal components of standardized waveforms from the training set. Three classes of behaviors could be identified from this classifier, including aversive gapes, neutral/appetitive MTMs, and no movement. All detected behavioral events were marked by a start and end time. For more details, see Baas-Thomas et al., in press; code for generating and running the classifier, as well as the final training dataset can be found at https://github.com/katzlabbrandeis/blech_emg_classifier.

#### Conversion of gape, MTM, and no movement events into timeseries frequency data

Behavioral events detected by the XGBoost classifier were converted into time-resolved binary vectors by discretizing the post-stimulus interval into uniformly spaced temporal bins of 3.5ms length. For each trial of each taste, bins overlapping with a detected behavioral event were assigned a value of 1, while all other bins were assigned 0. Binary vectors were then summed across trials to compute the proportion of trials containing the behavior at each time point for a single taste. Frequency traces were normalized by total taste trials and smoothed using a moving average filter (window size = 20 bins) prior to visualization. This amounts to a final smoothing window of 70ms, which falls well-below the 100-250ms cycle range for possible detected behaviors ranging from 4-10Hz (see next section).

#### Gape bout detection

Gapes occur in rhythmic bursts of 2-6 movements falling within a frequency range of 4-6Hz, or one gape every ∼166-250ms (Grill & Norgren, 1978a). The first gape bout is defined as the first series of a minimum of 3 consecutive gapes labelled by the XGBoost classifier with an inter-gape interval of 167-250ms following taste delivery (Grill & Norgren, 1978a) and an onset no later than 2000ms after stimulation (after animals have made consummatory decisions; Katz et al., 2001; Sadacca et al., 2016). Statistical comparisons between first gape bout onset distributions were performed using Mann-Whitney U tests.

#### Correlation of MTM frequency to taste palatability ranks

To determine if the MTM frequency for each taste is significantly correlated to hedonic rating, or palatability ranks, of tastes during quinine sessions, binned trial-wise binary vectors for MTMs post-stimulation (see *Conversion of gape, MTM, and no movement events into timeseries frequency data*) were summed for each trial. The resulting trial-wise MTM sum for all tastes were concatenated and a corresponding vector with the palatability rank assigned to each trial was created as follows: 1mM QHCl = 1, 0.1mM QHCl = 2, water = 3, 0.1M NaCl = 4. Trial-wise behavioral sums were correlated with palatability ranks using Spearman rank correlation. The distribution of quinine-session Spearman rank correlation coefficients (rho) across animals was compared to 0 (chance) using a one-sample t-test.

#### Quantification of gaping activity around CS delivery

To quantify the amount of gaping activity in response to NaCl and water before and after sacc CS was introduced into the session, gape occurrences were quantified around the last trial for NaCl and water before CS was introduced to the taste panel. Gapes for NaCl and water were summed within each of the 10 trials before and 15 trials after CS delivery (these analyses were only performed on trained and over-trained animals, which each had a minimum of 10 NaCl and water deliveries before the CS introduction, see **Table 1**). Trial-wise gape sums for NaCl and water trials were then concatenated and summed across sessions. The palatable taste gape sum pre and post-CS introduction was then normalized to the number of trials (a separate normalization factor was used for the pre vs. post-CS data because there were fewer pre-trials). To test for significant differences in gape counts before and after the CS, the mean number of NaCl and water gapes per trial pre vs. post-CS were independently quantified for each session and compared statistically using a paired-samples ttest.

#### Modelling sudden saccharin gape onset changes across trials

To quantify the abrupt transition to early sacc CS gaping across trials, sigmoid functions were fit to pooled trial data. First gape bout onset was plotted as a function of normalized trial number and fits were performed using a four-parameter logistic model (scipy.optimize.curve_fit, SciPy):

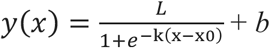

where (L) denotes the dynamic range of the response, (k) controls sigmoid steepness, (x0) corresponds to the inflection point (transition midpoint), and (b) represents baseline offset. Initial parameter estimates were initialized from the observed data range and optimization was performed with a maximum iteration limit of 10,000 function evaluations. The fitted inflection point was used as a measure of transition timing. A distribution of inflection points from individual session fits was used to determine the median transition time to ‘early’ gaping.

#### Electrophysiology

##### Single Unit Isolation

Single unit isolation spikes from electrophysiological recordings were sorted and analyzed off-line using an in-house Python-based pipeline (Mahmood et al., 2025). Putative single-neuron waveforms with >5:1 signal-to-noise ratio (median absolute deviation) were sorted using a semi-supervised algorithm: recorded voltage data were filtered between 300-3000Hz, grouped into potential clusters by a Gaussian Mixture Models (GMM) fit to multiple waveform features; clusters were then labeled and/or refined manually (to increase conservatism) by the experimenters.

##### Neural population changepoint estimation & behavioral alignment

GC ensemble taste responses have been repeatedly shown to involve sudden, coherent firing-rate transitions. Because these transitions typically involve firing-rate changes in ∼50% of the neurons in simultaneously recorded ensembles (Jones et al., 2007), and because they are sudden and stark (see below), they can be observed in single trials using any of a number of methods (see Mukherjee et al., 2019) and can be robustly inferred from ensembles as small as six neurons (Jones et al., 2007). Here, a multi-changepoint model written in the probabilistic programming language pymc3 (Salvatier et al., 2016) was used to determine the presence and latency of changes in ensemble responses. For a detailed explanation of the changepoint model used to identify ensemble transition times, see Mahmood et al. 2023 (code can be accessed online at https://github.com/abuzarmahmood/pytau/blob/development/pytau/examples/ Bayesian_Changepoint_Model.ipynb).

To align gape frequency with GC changepoint 2 (the onset of palatability-related firing; Sadacca et al., 2016; Mukherjee et al., 2019), a 4-state changepoint model was fit to the neural ensemble activity (Mahmood et al. 2023), outputting 3 changepoints per trial. Trials were aligned to changepoint 2, creating a peri-transition time matrix. In 400ms bins before and after the changepoint, categorical event outputs of the XGBoost classifier were converted into a Boolean array representation of gaping or no movement at each 1ms time point (1 assigned for timepoints in which the behavior was occurring and 0 for timepoints in which the behavior was not occurring). The gape or no movement Boolean arrays were then summed across trials and normalized by the total number of trials in the session before visualization.

##### Estimation of behavior suppression and peak after changepoint 2

To determine the latency to peak of QHCl gaping and trough of NaCl and water gaping after inferred neural changepoint 2, trial-wise gape Boolean vectors aligned to 400ms before and after the changepoint (see *Neural population changepoint estimation & behavioral alignment*) were averaged across trials to create a single session trace. Traces were smoothed using a one-dimensional Gaussian filter (scipy.ndimage.gaussian_filter1d) with σ = 2 bins to reduce high-frequency noise and stabilize latency estimates. Gaussian smoothing replaced each time point with a weighted average of neighboring bins according to a Gaussian kernel:

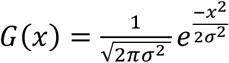

For QHCl, the latency of maximal behavioral expression was quantified as the time of the maximum value in the smoothed post-changepoint trace. For water and NaCl, the latency of behavioral suppression was quantified as the time of the minimum value in the smoothed post-changepoint trace. Peak and trough latencies were extracted separately for each session.

Latency distributions were visualized using bar plots with overlaid individual-session data points. Statistical comparisons between the high-QHCl peak latency distribution and each trough latency distribution were performed using a two-sided Mann–Whitney U test.

## Results

### Naïve rats demonstrate aversion to bitter quinine by producing long-latency gapes

Gaping is a well-characterized indicator of aversive behavior known to occur in response to both naturally bitter quinine and tastes rendered aversive through CTA learning (Grill & Norgren 1978a, Berridge et al., 1981; Pelchat et al., 1983; Berridge & Spector et al., 1988; Parker et al., 2003 & 2006). To investigate whether the form and neural underpinnings of these gapes differ, we implanted taste-naïve rats with electrodes in gustatory cortex (GC; A/P +1.4, M/L -5.0, D/V -4.7mm), electromyography (EMG) wires in the anterior digastric (jaw-opener) muscle, and an intraoral cannula (IOC) for taste delivery directly to the tongue.

After 5d of surgical recovery and 2d of habituation to receiving water deliveries through the IOC, animals underwent a three-day CTA protocol: the first day consisted of a single training session, during which the rats received randomly ordered 60μl aliquots of 5mM saccharin (sacc; the conditioned stimulus or CS) and water *via* IOC, followed by a subcutaneous injection of 0.6M LiCl (1% v/w) 30 min later; the second day consisted of a testing session, during which the rats received 60μl IOC deliveries of water, 0.1M NaCl, and the sacc CS; the third and final day consisted of a session in which they received 60μl IOC deliveries of water, 0.1M NaCl, and 1mM quinine hydrochloride (QHCl) called the “quinine session” (trained group, n = 7 animals; **Figure 1A**; **Table 1**). We used a strong concentration/dose of LiCl as an unconditioned stimulus to ensure that aversion was communicated *via* gaping to sacc CS in the testing session (Li et al., 2016). An additional, control cohort of rats (sham-trained group; n = 5 animals) received an identical protocol, but were injected with 0.6M NaCl (1% v/w) after the training session instead of LiCl (**Table 1**).

We isolated 3 classes of orofacial behaviors from EMG signals recorded in both cohorts by using a machine-learning classifier (XGBoost) trained on features extracted from manually labelled anterior digastric muscle EMG datasets, as done previously (Baas-Thomas et al. in press; **Figure 1B**). An advantage of the XGBoost classifier is that it detects not only aversive gapes (4-6 Hz), but also MTMs, a heterogenous class of behaviors encompassing both appetitive tongue protrusions (8-10 Hz) and neutral mouth movements (6.6-8 Hz; Grill & Norgren, 1978a), and low-amplitude segments of the signal in which the animal is not moving its mouth (called “no movement”). Thus, this classifier provides a detailed assessment of hedonically mixed behavioral responses on a single-trial level (Berridge et al., 1981; Berridge & Grill 1983) that we might expect from rats as they develop a CTA.

The algorithm detected few gapes to sacc in naïve rats (i.e., prior to training). Meanwhile, QHCl gaping was prominent in the quinine session. To validate our protocol, we went on to analyze the timing of QHCl gapes in rats without CTAs (i.e., sham-trained rats), which are known to have an average onset latency of between 0.8 and 1.0s following taste delivery (Li et al., 2013; Sadacca et al., 2016, Mukherjee at al., 2019); as predicted, the classifier reliably detected gaping responses to quinine with an approximate onset of >1000ms post-delivery (**Figure 1C**). These results confirm the effectiveness and accuracy of our oral behavior classifier.

### Conditioned aversion to saccharin is reflected in shorter-latency gaping bouts

To confirm that our protocol induced reliable CTAs, a subset of trained and sham-trained rats were given 30min home cage access to 15ml of the CS and, subsequently, water. We did not measure CS consumption in the training session, and therefore did not perform within-animal comparisons of the amount of CS consumed pre- and post-training. Instead, we compared sacc CS consumption between the trained and sham-trained groups, a protocol that allowed us to avoid: 1) attenuating the eventual CTA by presenting the CS in two difference contexts (i.e. *via* bottle vs. IOC) across training and testing (Parker, 2006) and 2) extinguishing the CTA by giving animals access to CS via a bottle after training (Rosas & Bouton, 1996; Nolan et al., 1997; Moran & Katz, 2014).

This analysis revealed that sham-trained rats drank significantly more CS (8.1 +/- 1.40 ml) than trained rats (1.25 +/- 0.34 ml; t = -4.75, p = 0.033; purple bars in **Figure 2A**). The amount of water consumption (blue bars) did not differ significantly between groups (sham-trained: 3.03 +/- 1.89 ml, trained: 4.25 +/- 1.59 ml; t = 0.49, p = 0.65), indicating that satiation level likely did not contribute to sacc consumption differences.

**Figure 2.**
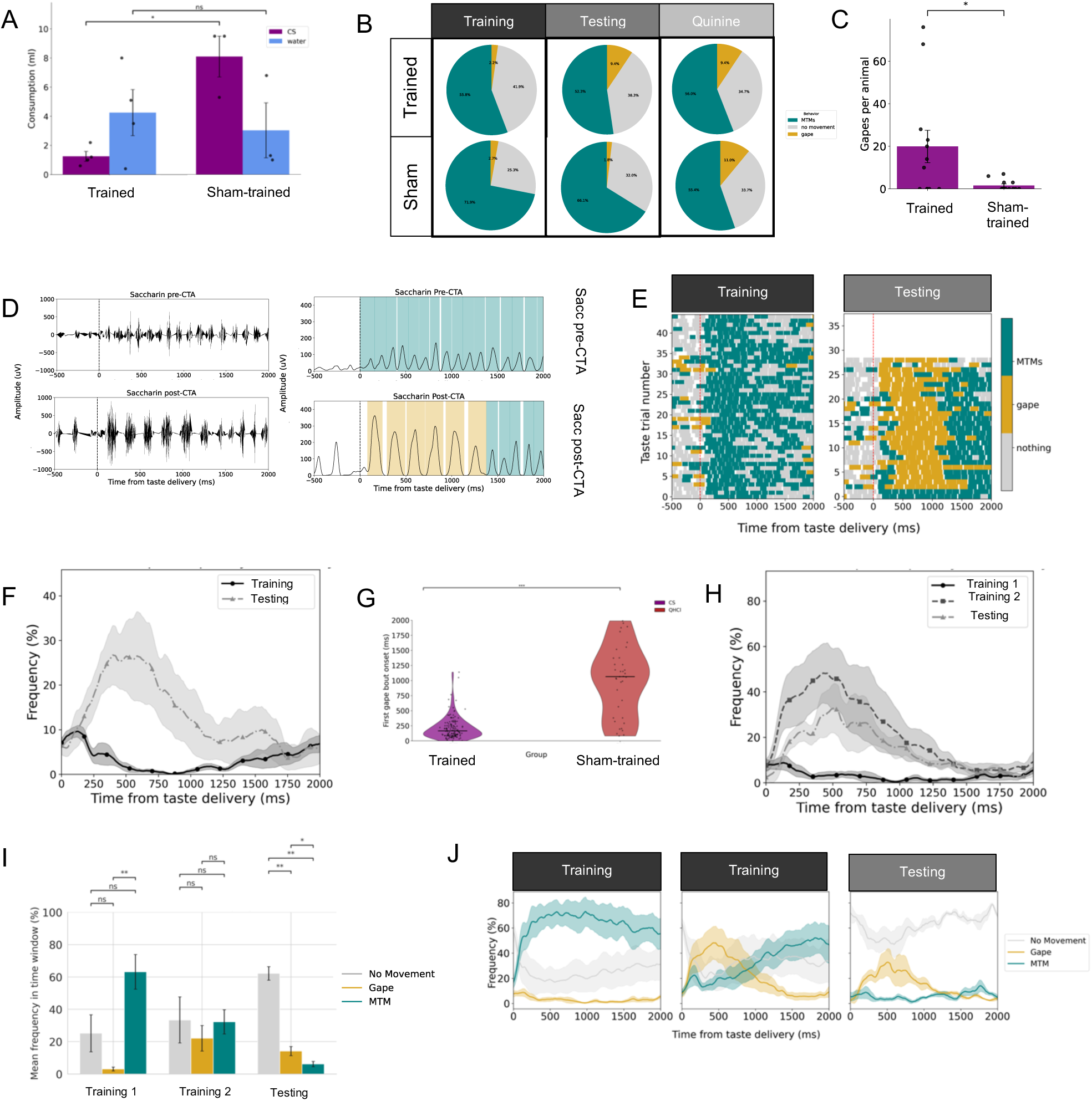
CTA-induced gapes have a shorter latency than naïve gapes. **A)** Average sacc CS consumption in 30 min bottle test is significantly less in trained than sham-trained animals (t = - 4.75, p = 0.033). Error bars represent SEM. **B)** Relative frequencies of XGBoost classifier-labelled behaviors in trained and sham-trained groups during training, testing, and quinine sessions. **C)** Average number of gapes during testing session sacc CS trials across animals is significantly higher in trained than sham-trained group (U = 40, p= 0.047). Error bars represent SEM. **D)** Example filtered (left) and enveloped (right) EMG signals recorded during two example sacc trials, pre-CTA (top) and post-CTA (bottom). Gape and MTM waveforms are highlighted in yellow and teal, respectively. **E)** Example raster plot of behaviors detected by classifier in response to all sacc trials during the training and testing session in one animal. **F)** Average frequency of gapes in response to sacc across trained animals (n = 7 animals) during training session (black, dot) and testing session (light gray, dash). Shaded area represents SEM. **G)** Distribution of first gape bout onset for sacc CS in trained group (n = 7 animals) and QHCl in sham-trained group (n = 5 animals). Horizontal black lines represent median. **H)** Average frequency of gaping in response to sacc CS across animals in over-trained group (n = 4 animals) before training (black, dot), after one training session (medium gray, square), and after two training sessions on testing day (light gray, dash). Shaded area represents SEM. **I)** Mean frequency of each behavior in response to sacc CS across the CTA paradigm in over-trained animals reveals significantly more ‘no movement’ detections during testing (no movement vs. gapes: t = 6.97, p = 0.0060; no movement vs. MTMs: t = 9.76, p = 0.0023). **J)** Average frequency of the 3 behavior classes in response to sacc CS across animals in TT group on each day (n = 4 animals). Shaded area represents SEM.

Learned aversions to sacc were confirmed in an analysis of gaping. Rats in the trained group produced more gapes in the testing session than in the training session, but rats in the sham-trained group did not, an expected outcome of successful conditioning. While MTMs constituted the majority of behaviors produced by both groups (gapes were comparatively seldom), the incidence of gapes made by trained rats more than doubled from the training to testing sessions, a result that was not observed for sham-trained rats (**Figure 2B**). A direct between-group comparison of the mean number of CS gapes produced per animal during the testing session revealed significantly more CS gaping in the trained group (U = 40, p= 0.047; **Figure 2C**). Clearly, trained rats found sacc aversive, whereas sham-trained rats did not; meanwhile, both groups found QHCl aversive.

It was immediately clear, however, that CTA-related sacc gaping differed in an important way from naïve QHCl gaping. **Figure 2D** shows a pair of trials from an example rat (raw EMG to the left, response envelope with the results of the machine-learning classifier to the right): in the training session (top row), this rat responded to sacc with low amplitude, high frequency MTMs; after a single training session, it began gaping to the CS. Note, however, that this gaping began at a very short latency—almost immediately following CS delivery. This example trial was representative of the majority of trials in the session (**Figure 2E**), and also of the entire testing-session dataset for trained rats, which showed gape frequency rising almost immediately, peaking 400ms after sacc delivery, and returning to baseline likelihood by approximately 1100ms after taste delivery (**Figure 2F**). A direct comparison of gape onset latencies to sacc CS in trained rats and QHCl in sham-trained rats (see Methods for explanation of how gape bouts are identified) confirms the significant difference between the two “types” of gaping (t = 13.40, p = 2.50e^-28^; **Figure 2G**). In both cases, the proffered tastes were deemed aversive, but this aversion was expressed with very different latencies.

It’s worth noting that in the case of extreme CTA, gaping is not necessarily the behavior actuating the aversive reaction to a proffered taste, a fact demonstrated in a third cohort of rats given a sequence of two identical training sessions separated by 24 hr (over-trained group, n = 4 animals). The peak of CS gape likelihood in these highly trained rats was not higher than that for rats receiving only one training day; in fact, gaping likelihood declined slightly between the second training session (48.25 +/- 13.09 trials, 428ms) and the subsequent testing (32.792 +/-10.91 trials, 509ms; **Figure 2H**). Whereas MTMs are the dominant oral behavior in the initial training session, and gapes increased in training session 2, after 2 training sessions the most prominent behavior was no behavior at all—strong aversion is essentially reflected in slack lack of oral response (no movement vs. gapes: t = 6.97, p = 0.0060; no movement vs. MTMs: t = 9.76, p = 0.0023; **Figure 2I**). Notably, the latency of this aversion-related cessation of movement was early, and similar to that of gaping (**Figure 2J**). These results demonstrate that while gaping reflects taste aversiveness, that relationship is not linear; gaping most consistently occurs when a rat is experiencing an intermediate level of aversion.

### Post-CTA early gaping generalizes to naturally aversive tastes and even palatable tastes

The above analyses suggest the existence of two “types” of gapes that are distinguished by their onset and offset latencies; **Figures 2I** & **2J** add further evidence that the decision that CS-sacc is aversive is made very quickly—quicker than the decision regarding QHCl. Such temporal differences in behaviors further imply differences in underlying neural circuitry, and it is tempting to attribute these differences to training—that is, to conclude that aversion to naturally aversive tastes arises out of the function of a relatively slow circuit, whereas aversion to the CS of CTA training arises out of a more quick-processing circuit. Analogous differences between naïve and learned behaviors have been attributed to learning in other paradigms (McCormick et al., 1981, 1982, & 1984; Steinmetz et al., 1987 & 1992).

The very fact that our analyses were performed on two distinct groups of rats—one of which received (one or two sessions of) CTA training, and one of which was sham-trained—renders this interpretation tentative, however. A stronger test of the above hypothesis requires the examination of gaping to QHCl in trained rats. We performed this test, first comparing quinine-session gapes in trained and sham-trained rats. This analysis revealed that our initial hypothesis was incorrect, in that CTA training appears to induce a general change in gaping even to naturally aversive tastes: the median gape latency following QHCl administration, which was 1065 ms in sham-trained rats, was only 424 ms in rats that received a CTA training session (U of difference from sham-trained = 690.6, p = 1.4e^-4^; **Figure 3A**); similarly, over-trained rats receiving two training sessions also gaped earlier to QHCl than sham-trained animals (mean onset = 639 ms; U = 604.0, p = 0.0042).

**Figure 3.**
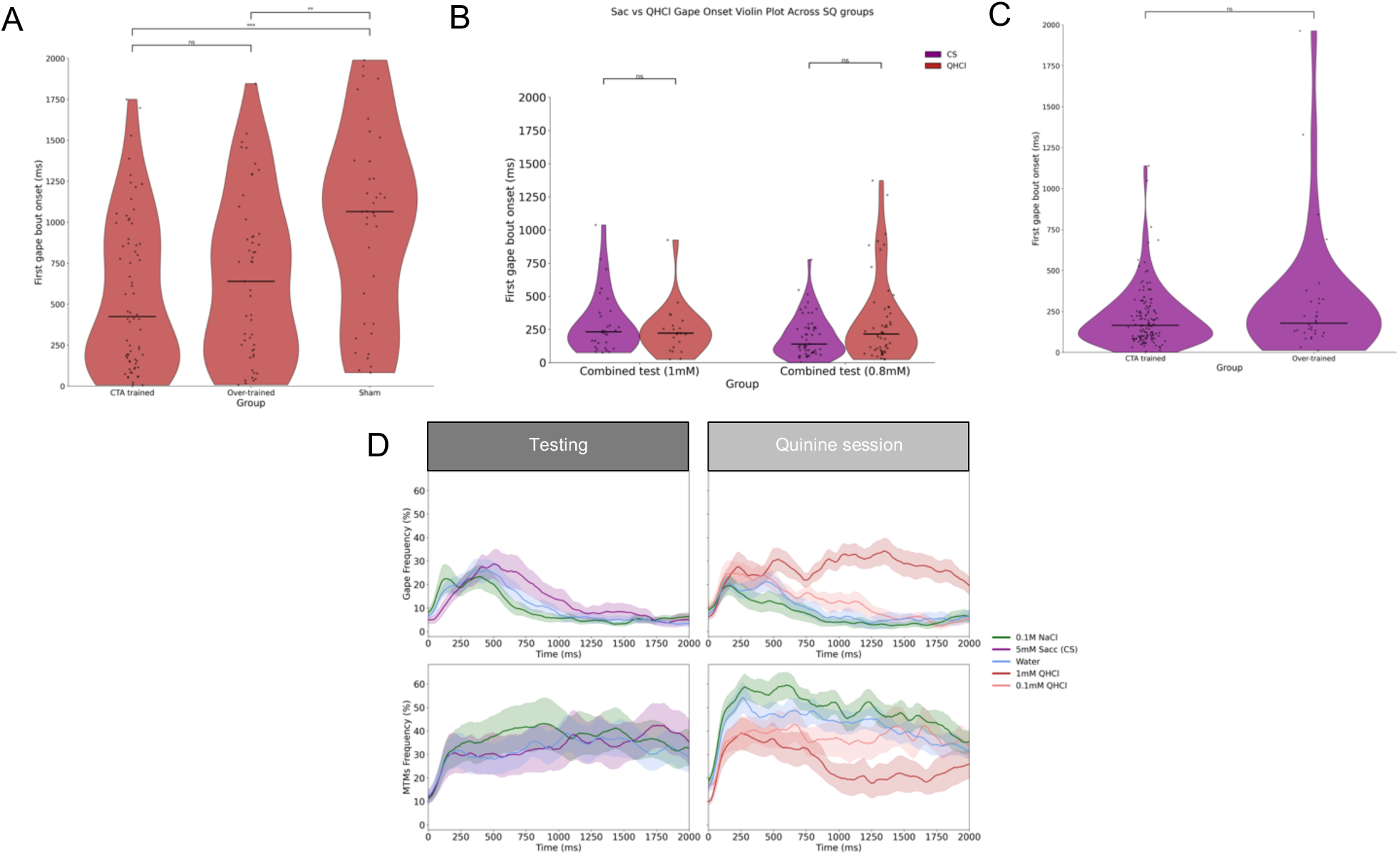
Post-CTA gaping generalizes to aversive and palatable tastes. **A)** Distributions of QHCl first gape bout onset in trained, over-trained, and sham-trained animals. **B)** Distributions of gape bout onset for sacc CS and QHCl in each group that received a “combined test” of sacc CS and QHCl in the same session post-CTA training. **C)** Distributions of gape bout onset for sacc CS in trained and over-trained groups. **D)** Gape and MTM frequency averaged across trials and across all trained and over-trained animals combined (n =11 animals) during testing (left column) and quinine sessions (right column). MTM frequency is significantly correlated to taste palatability ranks during the quinine session (mean rho = 0.221 +/- 0.096; rho vs. 0: p = 0.045). All behavioral frequency and onset analyses were restricted to 2000ms following taste delivery.

Two additional details of **Figure 3A** are worth noting: 1) the lack of difference between QHCl gaping latency in trained and over-trained rats suggests that CTA intensity does not itself determine QHCl gaping latency; and 2) the evidence of bimodality in the distribution of latencies in QHCl gaping among CTA-trained rats suggests further complexities in the process(es) underlying aversion decisions. Both of these points will be discussed more below.

To more rigorously test the effect of CTA experience on QHCl responses, we ran additional groups of rats that received both CS sacc and QHCl (as well as water) in the testing session (“combined test” group; **Table 1**). A subset of these rats (n = 2) were offered the same concentration of QHCl as the other three cohorts (1mM), and others (n = 4) consumed a lower concentration of 0.8mM. Analysis of these sessions failed to reveal any significant differences in the gaping onsets to sacc CS (1mM: 232 ms; 0.8mM: 140ms) and QHCl (1mM: U = 358, p = 0.55; 0.8mM: U = 1206.5, p = 0.163; **Figure 3B**), indicating that prior CTA training experience hastened gaping responses to naturally aversive QHCl, suggesting that the mechanism driving aversive responses to a sacc CS may “co-opt” the driving of natural aversion.

Of course, another possible explanation for the above results is that a learned aversion to sacc is extreme, and that CTA experience also increases the aversiveness of subsequently experienced QHCl; gaping onset latency is known to shift earlier with higher aversiveness (Grill & Norgren, 1978a). We consider this explanation unlikely, however, because neither 1mM QHCl gaping (**Figure 3A**) nor sacc CS gaping was hastened by increased CTA training (one vs two training sessions, U = 1876.0, p = 0.30; **Figure 3C**). Thus, we provisionally conclude that different underlying mechanisms appear to underlie the two distinct latencies of gapes, and that the mechanism triggering early aversion evaluation may “overpower” normal aversive response mechanisms.

Further investigation revealed more differences between early and late evaluation, showing that CTA training drastically alters how the animal responds to even normally palatable stimuli. The testing session taste panel included, in addition to sacc, both water and a normally preferred (Sadacca et al., 2012) concentration of NaCl (0.1M, see **Figure 1A**), a fact that allowed us to observe that CTA-trained rats gaped to salt and water during both the testing and quinine sessions (**Figure 3D**). Sham-trained rats produced no significant gapes to these palatable tastes (data not shown). This salt and water gaping had almost identical temporal dynamics to CS sacc gaping, rising early and then vanishing at approximately 1000ms post-delivery; gaping to these tastants was clearly distinguishable from that to strong QHCl, in that the likelihood of gapes to the naturally aversive stimulus persisted (and in fact increased, as predicted) >1000ms after taste delivery.

This generalized gaping occurred despite the fact that MTM frequency in the same session (**Figure 3D**, bottom right), another reliable indicator of taste palatability (Grill & Norgren 1978a; Travers & Norgren 1986), was significantly correlated with natural taste palatability ranks (mean rho = 0.221 +/- 0.096; rho vs. 0: p = 0.045). These results demonstrate that CTA-trained rats are fully capable of distinguishing palatability, but they apparently make early, palatability-independent aversiveness judgments.

### Early, palatability-independent aversiveness judgments appear in a sudden state change, driven by administration of CS sacc

The above work suggests that there are 2 distinct drivers of aversive responding to fluids in the mouth, but that the distinction isn’t as simple as “gaping to a substance that is naturally aversive versus gaping to a taste that the rat learns to dislike.” The distinction seems to have more to do with the animal—“gaping by a rat that hasn’t yet received a CTA vs gaping by one that has.” That is, it appears that early gaping represents some sort of state switch in which a rat that has been put through unpleasant taste learning tends toward responding to all tastes with aversion.

Further analysis supports this conclusion, revealing that early, generalized (i.e., not taste-specific) aversion responses begin happening suddenly within sessions—an onset that appears driven by presentation of sacc CS trials themselves. Water and NaCl gaping in rats that received one training session, for instance, appears suddenly upon introduction of sacc CS (**Figure 4A & B**—an observation that we could make because we always began post-CTA sessions with a brief series of nominally innocuous tastes). After CS introduction, animals gaped significantly more per trial (t = -2.43, p = 0.033, n = 2 tastes per 6 sessions; **Figure 4B**, inset).

**Figure 4.**
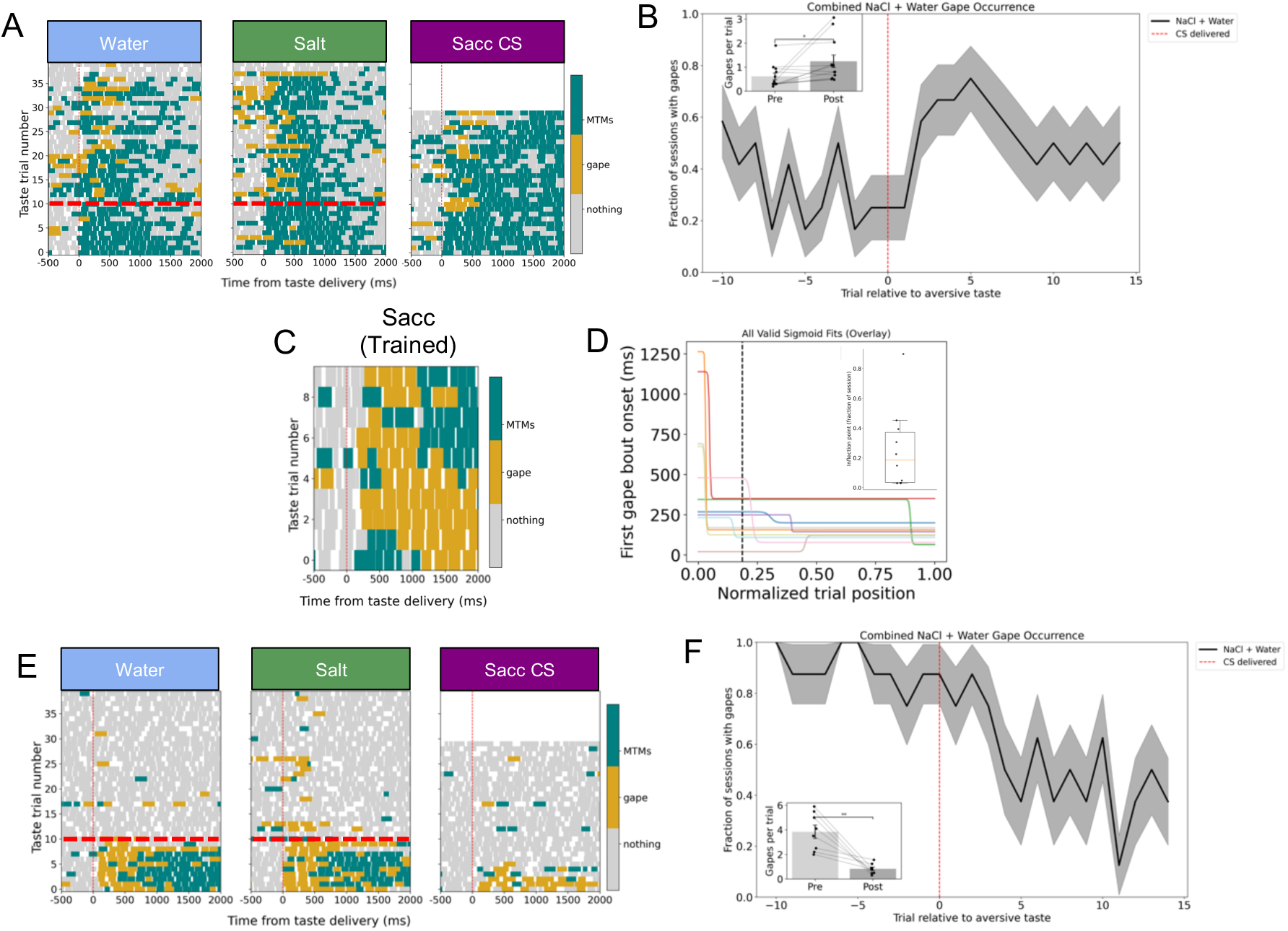
Palatability-independent aversiveness judgments appear in a sudden state change driven by administration of CS sacc. **A)** Example behavior raster plots for 3 tastes given during testing in an example trained animal; red horizontal dashed line represents last trial before onset of sacc CS deliveries. **B)** Fraction of testing sessions in which gapes were detected during both NaCl and water trials aligned to the onset of aversive taste deliveries. Inset: mean gape count per NaCl and water trial before and after CS introduction during the testing session in trained animals (t = -2.43, p = 0.033, n = 2 tastes per 6 sessions). **C)** Example behavior raster plot for the first 10 deliveries of sacc CS given during an example testing session for a trained animal shows a sudden shift to early gaping after trial 2. **D)** Successful sigmoid fits for first gape bout onset as a function of normalized sacc CS trial for 10/15 individual testing sessions taken from all three CTA-trained cohorts overlayed. Black vertical dashed line represents the median inflection point across all successful fits (0.186), the distribution for which is plotted in the inset. **E)** Example behavior raster plots for 3 tastes given during the testing session in an example over-trained animal; red horizontal dashed line represents last trial before onset of sacc CS deliveries. **F)** Fraction of testing sessions in which over-trained animals gaped to NaCl and water aligned to the onset of sacc CS deliveries. Inset: mean gape count per NaCl and water trial before and after aversive taste introduction during the testing session in over-trained animals (t = 4.66, p = 0.0023, n = 2 tastes per 4 sessions). All behavioral frequency and onset analyses were restricted to 2000ms following taste delivery.

And just as water and NaCl behavioral responses changed after 1-3 sacc CS trials, so did the sacc CS responses themselves. **Figure 4C** shows an example session of sacc CS responses, in which it can be seen that, for the first few trials, gaping onset to sacc CS was late, with a latency similar to that of QHCl gaping in sham-trained rats (see **Figure 1C**). This shift from late to early sacc CS gaping occurred in most rats, but as it did not occur on the same trial for every rat, we plotted gape bout onset as a function of normalized trial position and fit a sigmoid to these data for each testing session from individual animals. We were able to achieve successful sigmoid fits for 10/15 (66.7%) sessions (**Figure 4D**) and determined the median inflection point, or median time at which the rats exhibited a sudden hastening of sacc CS gape onset, to be about 18.6% of the way through the session’s sacc CS trials (ranging from trial #3-6; **Figure 4D**, inset).

The same sudden state-change could be observed in over-trained rats, who entered the testing session (following their two training sessions) already prepared to find water and NaCl aversive at short latencies: in these rats, the introduction of sacc CS precipitated a dramatic further increase in the perceived general aversiveness of fluid delivery, in the form of a sudden reduction in the incidence of orofacial behavior in response to water and NaCl (**Figure 4E & F**); after CS introduction in this context, animals gaped significantly less (t = 4.66, p = 0.0023, n = 2 tastes per 4 sessions; **Figure 4G**, inset). In these over-trained animals, inflection points for successful sigmoid fits (3/4 of the animals) reveal the state transitions happened very early in the session—after only 3% of the sacc CS trials.

### Two classes of gapes have different relationships to GC activity

The above analyses characterize two types of gapes: ‘late gapes’ that occur in response to unpalatable tastes, >1000ms post-delivery; and ‘early gapes’ to all delivered stimuli that typically appear within ∼200ms of stimulus delivery, in trials occurring after introduction of CS sacc to the taste battery. Given that the two types of aversion decision reflected in these distinct gapes specifically differ in both their taste dependence and timing, it is reasonable to predict that they have different relationships to the known taste-processing dynamics in GC. Late gaping has already been shown to represent the culmination of taste palatability processing; we hypothesized that early gapes, which are not taste-specific, would precede this firing-rate transition (the obvious alternative hypothesis being that GC response dynamics are altered with the transition to the early-gaping state).

To test this hypothesis, we analyzed ensemble single-neuron activity from GC (**Figure 5A**) that had been recorded during all days of the CTA paradigm. **Figure 5B** shows a representative set of PSTHs from a single GC neuron during the quinine session (the inset shows the average waveform for this neuron). This neuron, like many recorded in GC during tasting (Katz et al., 2001, Jones et al., 2007, Sadacca et al., 2016), progressed through 3 firing-rate states once a taste hit the tongue—first processing stimulus detection, then chemical identity, and finally palatability (note the changes in response ordering at ∼200ms and ∼1000ms after stimulus delivery). We used a Gaussian Changepoint Model developed to detect the time points that best describe when the population of simultaneously-recorded neurons changed their firing rates in single trials. The model returned the expected 3 changepoint times, corresponding to the transition from detection to identity processing (CP1), the transition from identity to palatability processing (CP2), and the end of palatability processing (CP3); CP2 is the changepoint that has been repeatedly associated with the decision that a taste is aversive (“palatability-onset” in ensemble responses; Sadacca et al., 2016; Mukherjee et al., 2019; Mahmood et al., 2026). We hypothesized that salt and water gapes would precede palatability-onset, and that the likelihood of gaping to QHCl would increase at the start of palatability-onset (Sadacca et al., 2016; Mukherjee et al., 2019; Baas-Thomas et al., in press; see also **Figure 3**)—given the complex impact of CTA, we made no specific prediction for CS-sacc (but see below and Discussion).

**Figure 5.**
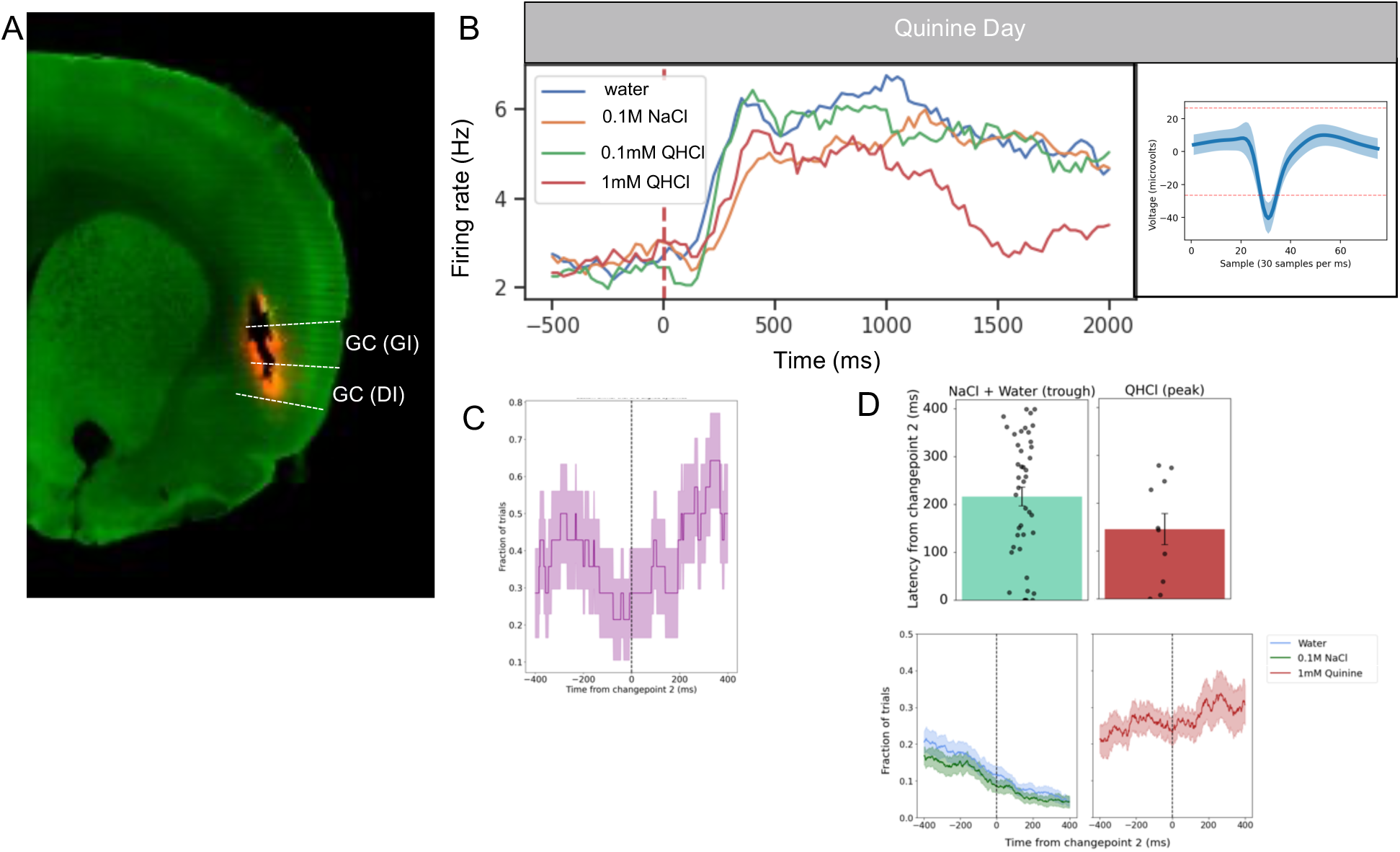
Two gape classes have different relationships to GC activity. **A)** Coronal slice from 1 rat. The track is an electrode bundle lesion, labeled with Vybrant Dil cell marker, with the end of the lesion marking the final resting location of the wires. **B)** The average waveform of one representative putative neuron during a quinine session; light blue shading represents the standard deviation of the noise in the waveform (right) and corresponding PSTHs (left). **C)** Fraction of trials with sacc CS gaping detected within 400ms of CP2, averaged across the first trial in which a gape bout was detected for the session in trained, over-trained, and combined test animals (n = 1 trial per 16 sessions; 2 animals lacking true bouts were excluded). **D)** Top: QHCl gaping peaks at the same time that NaCl and water gaping reaches a minimum after CP2 (QHCl peak vs. NaCl + water trough: t = -1.59, p = 0.118). Bottom: fraction of trials NaCl and water combined and QHCl trials with gaping detected within 400ms of CP2, averaged across all trained and over-trained animals. GC, gustatory cortex; GI, granular layer; DI, dysgranular layer; CP2, GC neural changepoint 2; CP1, GC neural changepoint 1.

The results of this analysis confirmed our hypothesis. Across the entire usable dataset, sacc CS gaping was plentiful before palatability-onset, and increased again after palatability-onset (**Figure 5C**). The first mode in **Figure 5C** reflected the fact that some sacc CS gaping declined before CP2, and the later mode reflected the fact that sacc CS gaping frequency increased at palatability-onset; therefore, sacc CS gaping can take on the form of both ‘early’ non-taste specific gaping as well as later palatability-associated gaping. NaCl and water gaping, on the other hand, specifically declines as palatability-onset approaches (**Figure 5D**, bottom left panel). Further examination of this phenomenon confirmed that the timing of the return to statistical baseline of water and NaCl gaping (average: 217ms after palatability-onset; **Figure 5D**, top left panel), which was not significantly different from the time that QHCl gaping peaked between 100 and 200ms following palatability-onset (t = -1.59, p = 0.118; **Figure 5D**, comparison of distributions in top panel).

In summary, our data reveal that the processing of tastes, which normally culminates in rats recognizing QHCl to be aversive, also culminates in rats in a “treat everything as aversive” state realizing that there is no need to reject NaCl or water.

## Discussion

### The interplay of two mechanisms whereby a fluid in the mouth may be deemed aversive

Here, we have provided evidence for two processes whereby an oral fluid may be deemed aversive (as indexed by gaping)—one that is driven by the sudden GC transition into palatability-related activity, as described by previous studies (Sadacca et al., 2016; Mukherjee et al., 2019), and a non-specific one with a significantly shorter latency that is squelched by that same transition. Rather than there being a single system involved in determining whether a fluid in the mouth is aversive, it appears that there are multiple such systems, differing in both their speed of operation and relationship to taste processing. We will discuss each of these in turn.

We observed “normal” late gaping to QHCl, but in our paradigm, the onset of that gaping was hastened by previous CTA experience. In rats experiencing QHCl concomitantly with the sacc CS after CTA training, the latency of gaping to QHCl is now indistinguishable from that of the CS, and is hundreds of milliseconds earlier than naïve gaping to QHCl that is 30x more concentrated (Travers & Norgren, 1986). In that situation, we can still see the function of both aversion-decision mechanisms, in that both QHCl and newly-aversive sacc can trigger gaping both early and late.

Note that further CTA training, which renders the sacc CS even more aversive (Berridge et al., 1981; Pelchat et al., 1983; Parker, 1988a), largely does not cause the rats to gape; once the animal has exceeded a certain threshold of aversive stimulation within a session, the actuation of the aversion decision appears to involve not oral behaviors but full-body aversive behaviors including chin rubbing, paw flailing, and rearing (data not shown, but see Berridge et al., 1981; Grill & Norgren, 1978a), and even an escape behavior like rearing or increased locomotion (Berridge & Grill, 1981 & 1983). We doubt, however, that this is a separate (third) mechanism of evaluating stimulus aversiveness, because it is similar to early gaping in its timing and generalization (and its state-dependence, see below).

### Early judgments of aversiveness emerge out of a change in internal state

In our hands, these early, generalized aversion judgments do not appear out of nowhere—they are not present in the initial responses to any tastes, even including CS-sacc. Rather, the introduction into the flow of tasting trials of a few deliveries of highly aversive taste, or of repeated aversive taste deliveries, appears to trip the rat into a state in which early aversion judgments—be those judgments expressed with gaping or a total cessation of oral behavior—occur to all tastes. Simply put, the rat enters a state in which it is primed to find any fluid in the mouth immediately aversive.

At this point, the nature of the internal state underlying the sudden increase in aversive behavior across all taste stimuli remains unresolved. We are tempted to speculate that it is an anxiety-like state triggered by the stressful experience of being back in a context in which the rat had consumed tastes that proved noxious. It is known that anxiety-like states alter taste perception in humans (Grunberg et al., 1992; Zushi et al., 2023) and taste behavior in rodents (DeSousa et al., 1998), and can cause context-dependent generalization of fear responses as well (Berton et al., 1998; Hebb et al., 2003; Andreatta et al. 2020). Early gapes may be analogous to generalized fear responses during an anxiety-like internal state caused by repeated aversive stimulation that put the animal in a heightened state of vigilance.

Testing of this hypothesis awaits further experimentation, but we would argue that it helps resolve conflict between a great deal of taste work and a classic study showing that rats subjected to CTA may cease licking at a spout offering the CS within 250ms (Halpern & Tapper, 1971). It was noted that this early rejection decision was made with some degree of response generalization, suggesting that it might be made prior to completion of taste processing, and might in fact reflect a generalized response; while generalization was not as complete as that observed here, that fact may reflect a difference in behavioral readout, or more likely a lack of complete descent into the anxiety state—it is reasonable to expect that even small details of a tasting paradigm (perhaps the number of aversive tastes delivered in a row, for instance) will impact the likelihood of entering such a state, and the robustness of that state.

### Taste processing kicks off one aversion-judgment mechanism and inhibits the other

One reason (beyond the generality of the response) to consider early gaping independent of taste processing is the relative slowness of that processing: the chorda tympani, the cranial nerve carrying taste information from the anterior tongue to the nucleus of the solitary tract in the brainstem, reaches peak response discriminability between stimuli only 200ms post stimulation (Faull & Halpern, 1972), which is too late to account for the early gapes; GC taste responses recorded with IOC delivery to passive rats doesn’t even appear until after this time point (Katz et al., 2001; Jones et al., 2007), further supporting the argument that animals are not only capable of rejecting taste stimuli without be able to maximally discriminate between them, but that they must do so. Relatedly, it has been shown that gaping can be induced in rats lacking a cerebrum (Grill & Norgren 1978b).

Given that these early decisions are being made without substantial gustatory processing, it is reasonable to ask whether GC, which is involved in the driving of naïve taste decisions (Sadacca et al., 2016; Mukherjee et al., 2019; Baas-Thomas et al., in press), and is necessary for normal CTA acquisition (Grill & Norgren, 1978c; Roman & Reilly, 2007) and late expression of CTA by gaping (Grill & Norgren, 1978c; Kiefer & Orr, 1992; Grossman et al., 2008), plays any role in early aversion judgments. Our analyses suggest that it does: first it appears that the “quick-judgment state” is only entered after trained rats first produce sacc CS gapes that have an onset after the GC palatability epoch onset, suggesting that taste processing is vital on this longer time-scale; furthermore, GC palatability processing is still relevant when rats are making quick aversiveness judgments, in that early gaping (most notably to NaCl and water) is quenched by the same onset of the palatability epoch that drives naïve gaping, to be replaced by an increase in appetitive MTM frequency.

In summary, taste processing dynamics appear to proceed normally even when the rat is making snap aversiveness judgments; when complete, this processing even serves to “correct” the judgment, letting the rat know that palatable tastes are indeed non-aversive and halting rejection behavior.

## Notes

### Competing Interest Statement

The authors have declared no competing interest.

### Funding

- NIH NIDCD: 5R01DC007703-20

